# Becoming a better parent: mice learn sounds that improve a stereotyped maternal behavior

**DOI:** 10.1101/2019.12.19.883041

**Authors:** Alexander G. Dunlap, Cristina Besosa, Leila M. Pascual, Kelly K. Chong, Hasse Walum, Dorottya B. Kacsoh, Brenda B. Tankeu, Kai Lu, Robert C. Liu

## Abstract

While mothering is often instinctive and stereotyped in species-specific ways, evolution can favor genetically “open” behavior programs that allow experience to shape infant care. Among experience-dependent maternal behavioral mechanisms, sensory learning about infants has been hard to isolate from motivational changes arising from sensitization with infants. We developed a paradigm where sensory learning of an infant-associated cue improves a stereotypical maternal behavior in female mice. Mice instinctively employed a spatial memory-based strategy when engaged repetitively in a pup search and retrieval task. However, by playing a sound from a T-maze arm to signal where a pup will be delivered for retrieval, mice learned within 7 days and retained for at least 2 weeks the ability to use this specific cue to guide a more efficient search strategy. The motivation to retrieve pups also increased, but that alone did not sufficiently explain the shift in search strategy. Bilaterally silencing auditory cortical activity significantly impaired the new strategy without changing the motivation to retrieve pups. Finally, motherhood as compared to infant-care experience alone accelerated how quickly the new sensory-based strategy was acquired, suggesting a role for the maternal hormonal state. By rigorously establishing that newly formed sensory associations can improve the performance of a natural maternal behavior, this work facilitates future studies into the neurochemical and circuit mechanisms that mediate novel sensory learning in the maternal context, as well as more learning-based mechanisms of parental behavior in rodents.

## 1. Introduction

Parental care in mammals is critical for a species’ survival, and understanding the neural mechanisms that control maternal behaviors has broad relevance for behavioral and systems neuroscience [1, 2, 3]. However, research into maternal behavior has largely been dominated by questions about its onset and maintenance. Consequently, we understand more about how hormones act in certain brain areas to promote maternal motivation than we do about the motor or sensory contributions to actually performing maternal care and improving at it.

In the case of sensory learning contributions to maternal behaviors, studies in mice have demonstrated experience-dependent auditory cortical plasticity for pup ultrasonic vocalizations (USVs) [4, 5, 6], which elicit a search and retrieval behavior from mothers. Progress has been made into the biological mechanisms mediating these neural changes [7, 8, 9], but the functional significance of this sensory cortical processing and plasticity for *how* maternal behaviors are performed remains unclear. One hypothesis has been that these changes alter how animals use sensory cues to carry out maternal behavior. Indeed, extrapolating from non-social auditory associative learning paradigms [10, 11, 12], plasticity in the auditory cortex could help establish a maternal memory trace for the increased behavioral relevance of the sounds pups produce [13, 14]. That putative sensory association could then functionally improve the delivery of infant care.

We sought to rigorously determine whether auditory associative learning about pup-predictive sounds can indeed enable more efficient execution of maternal behaviors. In order to validate sensory learning as it occurs, we sought to associate a novel sound with the same, innate maternal retrieval that animals naturally perform after finding a pup. One challenge to quantifying these sensory-driven changes in maternal behavioral performance is dissociating them from non-specific changes in the motivation to act maternally. Here we describe a method for driving a measurable, sensory-guided change in a maternal behavior that works for both lactating mothers and virgin mice. We used pup retrieval on a T-maze [15, 16] while playing back a non-ethological, otherwise neutral sound from a speaker co-localized with the arm from which a pup can be retrieved on any given trial. We characterized an initial strategy for selecting which arm to enter at the choice point of the T-maze and how that strategy changes to reflect the formation of a sensory memory about the association between the pup and the predictive sound. We then captured a separate measure of maternal motivation over the course of training to understand its relationship to the degree of associative learning. We validated that auditory cortical activity is needed to express the learned association but not maternal motivation. Finally, we determined how maternal state influences both of these behavioral measures.

## 2. Materials and Methods

### 2.1. Animals

All animal procedures were approved by the Emory University Institutional Animal Care and Use Committee (IACUC). Animals were female CBA/CaJ mice placed on a 14h light / 10h dark reverse light cycle with ad libitum access to food and water. Animals were socially housed in single-sex cages, however the animals that were cannulated were singly housed after their surgery. For mice that were cannulated, surgeries were performed one week before habituation began on the T-maze. All sound pairing experiments on the T-maze were conducted on mice aged 10-16 weeks.

### 2.2. T-maze behavioral paradigm for pairing novel sound with pup retrieval

Behavioral studies were conducted inside an 8’-2” × 10’-6” double wall anechoic chamber (IAC, Bronx, NY) under dim red light. We constructed an elevated (30 cm from ground) T-maze with the following dimensions: 86 cm overall length from the nest to the choice point of the T-maze stem, 2-cm-high walls, 8 cm arm width, 11 cm square nest width with a 1 cm drop from the stem floor to the nest floor. Speakers were located 25 cm along the y-direction from the end of the arms at the top of the T-maze and 10 cm along the x-direction from the end of the top of the T-maze. The speakers were slightly angled and oriented so that they faced the decision point where the stem intersected the arms of the T-maze. Prior to each test day, the T-maze was wiped down with Virkon disinfectant and clean Alpha-Dri bedding was added.

We habituated the animals to the T-maze for 3 days. Habituation consisted of 10 minute sessions on the maze without pups during which the experimenter would stay inside the anechoic chamber, occasionally opening and closing the chamber door every 2 minutes. This would acclimate the mouse to the experimenter’s presence and fluctuations in background sound level, which would occur when the experimenter entered or exited the testing chamber at the beginning and end of the experiment.

Sound pairing began after habituation on the third day and continued in consecutive daily sessions. During these days, a subject mouse was initially placed on the T-maze with two pups present for 10 minutes. After this time, the pups were scattered on the T-maze to encourage traversal of the maze. The first sound pairing trial began the moment the subject mouse retrieved the last scattered pup back to the nest. Subsequent trials began when the subject mouse retrieved the pup from the previous trial back to the nest. Mice that stopped retrieving for a whole session were discontinued from further pairing. At the start of each trial, the sound cue was streamed from one of the two speakers (Fig. 1). The speaker the sound was delivered from was chosen to always coincide with the arm where a pup would be delivered. Once the subject mouse left the nest and entered this arm (Fig. 1IV), sound playback was turned off until the pup was retrieved back to the nest.

**Figure 1:**
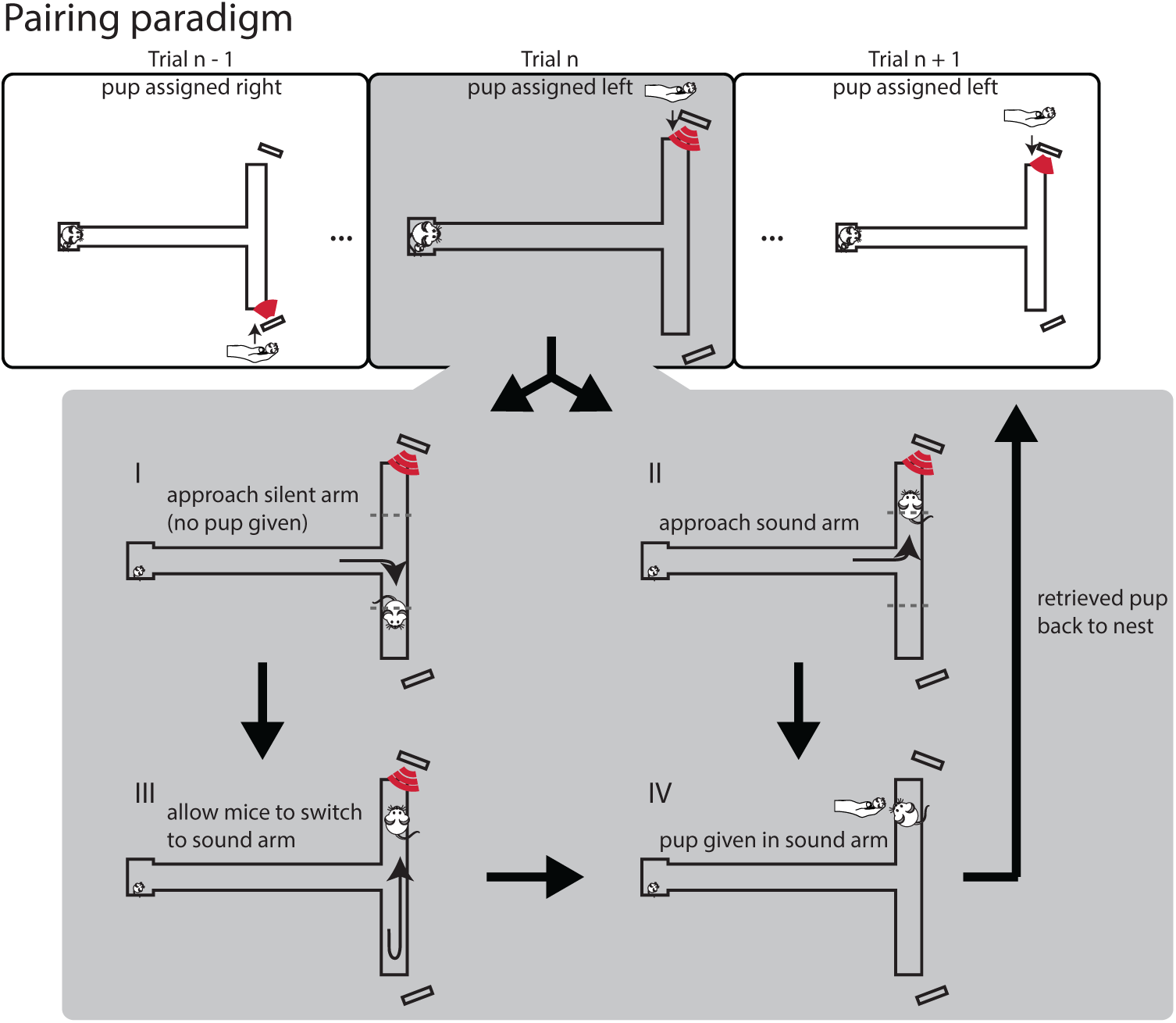
Retrieval paradigm and behavioral scoring. Each trial started by playing the auditory cue from one of the two randomly chosen speakers when the mouse was in the nest. (I & II) Once the mouse left the nest and its center of mass crossed a threshold within one of the two arms, we recorded the latency (measured as the time to cross threshold relative to the time the trial started) and the side that was chosen. (III) If the mouse entered the unpaired arm, then she was allowed to alternate to the other arm without punishment. (IV) Once the mouse found the paired arm she was manually given a pup, which she retrieved back to the nest to initiate the next trial. Before the mouse returned with the retrieved pup, the pup that had been left in the nest was removed and held outside the maze to be used in the next trial. Pairing lasted for 50 minutes or until the mouse completed 100 trials, whichever occurred first.

The side where a pup would be delivered was generated pseudo-randomly with probability 0.5. However, if the same side occurred consecutively for three trials, then the next side was forced to alternate. A red LED, elevated 30 cm above the maze and blocked from the subject’s direct view, indicated to the experimenter to which side to deliver a pup. Once the mouse found the correct arm, she was given a pup, which she retrieved back to the nest, initiating the next trial. Before the mouse returned with the retrieved pup, the pup that had been left in the nest was removed by the experimenter and held outside the maze to be used in the next trial. Training lasted for 50 minutes or until the mouse completed 100 trials, whichever occurred first.

### 2.3. Y-maze behavioral paradigm for testing sound memory

Behavioral studies were conducted inside a double wall anechoic chamber (IAC, Bronx, NY) under dim red light. We modified a rat-sized cage by adding a central dividing wall extending from one end of the cage to a third of the length of the cage’s long dimension. We then secured two vertically oriented plastic tubes against the side walls to create symmetric openings to the divided chambers from a nesting area at the other end of the cage. Holes were drilled into the rat cage’s wall at the end of each of the divided chambers to allow speakers placed behind the walls to broadcast into the cage. Following the same procedures as for the T-maze, naive mice could learn to use a sound to guide entry into a target chamber to retrieve a pup.

### 2.4. Sound stimulus

The sound stimuli used for the experiments presented in this paper were designed to be models of a previous recording of hand tapping on the T-maze (same amplitude modulation) but with a center frequency and bandwidth under parametric control. A vector of Gaussian white noise was bandpass-filtered (center frequency = 40 kHz, bandwidth = 33 kHz) and multiplied element-wise in time by the amplitude envelope of the hand tapping recording to create the final signal. From this final signal, two 20 second subsets were extracted to create “sound examples 1” and “2.” During playback the 20 second stimuli were streamed in a constant loop.

For the experiments presented in sections 3.1 - 3.3 sound example 1 was used for pairing on days 1 - 7, and example 2 was used on day 8. The purpose of the second sound example was to avoid pseudo-replication and verify that after training, mice would approach sounds with similar acoustic properties instead of a specific feature only in the recording of the training stimulus. Only sound example 1 was used in the experiments described in Sec. 3.5 - 3.6. For experiments described in Sec. 3.4, we used a 5 Hz sinusoidal amplitude-modulated noise (center frequency = 40 kHz, bandwidth = 33 kHz) with 100% modulation depth as the target stream. For experiments with a distractor sound stream, we used a frequency-modulated (FM) tone sweeping over the same frequency range as the non-target sound (from 30kHz to 50kHz). The FM tone was 200 ms long and generated with the following equation: *FM* = sin [2*π ×* (30000 + 50000 × (1 − exp −*t/*0.9))]. The FM tone was presented in a stream at the rate of 3 tones per second with a random gap between tones, to avoid rhythmic stimulation. The random gap had a mean of 1.333 s and was jittered with equal probability within a 40 ms window. The distractor sound and the target sound were presented simultaneously from the two speakers.

Sound stimuli were delivered by a TDT System 3 RX6 module, allowing the user to control trial initiation in real time via a custom made remote located inside the training room. The sound was played from Pioneer speakers (model PT-R4). For a given trial, the intensity of the sound at the decision point of the T-maze was fixed, while across trials it was roved over a range of 13 dB.

### 2.5. Analysis of approach behavior

Video of the behavior was recorded overhead using a Panasonic color cctv camera (model WV-CP284) mounted to the ceiling and connected to a PC located outside the anechoic chamber, which was running TopScan video tracking software. Custom Python and MATLAB code was used to identify when each trial started, when the subject mouse entered an arm, and which arm side was chosen. Trials were visually inspected to verify accuracy.

Once the mouse left the nest and crossed a threshold within one of the two arms, we recorded the latency (measured as the time to cross threshold relative to the time the trial started) and the side that was chosen. Latency values for each mouse were summarized over days as the reciprocal of their mean (reciprocal latency). Trials were scored according to both location- and sound-based strategies. For the location-based strategy, if the mouse did or did not choose to first enter the arm from which she received a pup on the previous trial, the trial was scored as correct or wrong, respectively. For the sound-based strategy, if the mouse did or did not choose to first enter the arm from which sound was playing on the current trial, the trial was scored as correct or wrong, respectively. The scores for each mouse according to a given strategy were summarized for each day’s session as percent correct.

### 2.6. Statistics

Individual mice were classified as having learned the sound-based strategy (Learner) if the sound-based strategy scores pooled over two consecutive days of pairing were significantly greater than chance (one-sample proportion test). For those experiments that did not use cannulated animals (Sec. 3.2, 3.3, & 3.6), trials were pooled over days 6 & 7. For the experiment that did use cannulated animals (Sec. 3.5), pairing occurred until animals reached the criteria for being a “Learner” or were removed from the study because they stopped retrieving (Sec. 2.2).

To account for the fact that we had repeated measures for scores (Fig. 4A, 7B-C, & 8C-D) and trial latencies (Fig. 4B, 5B-C, 7D, & 8A) within each animal, we used generalized linear mixed models to test for differences between groups. When comparing binary trial scores or continuous latency values, we fit our models using the binomial or the gamma family distributions respectively. These analyses were performed in R 3.4.1 using the lme4 package [17, 18].

For all statistical tests, significance was identified as *p* < .05.

### 2.7. Cannulation of auditory cortex

At least 1 week before the first day of habituation, mice in our muscimol study were bilaterally cannulated into auditory cortex (bregma = −3.04 mm, midline = +/-3.75 mm, z = −0.5 mm from the cortical surface). Cannulae were purchased from Plastics One (C315GS-4/SPC guide-cannula, stainless-steel, 26 gauge, 4mm pedestal size, cut length 4mm below pedestal). Mice were implanted with the cannulae under Isoflurane (2-5% with O_2_) after they showed no reflexive response to a light toe pinch. Once implanted, cannulae were covered with a dummy cap (Plastics One, C315DCS-4/SPC). During recovery, in order to prevent damage to the implanted cannulae, food was delivered through a side hanging hopper and water was delivered through a liquid diet feeding tube (Bio-Serv model 9019).

### 2.8. Drug infusion

For infusions, solutions of muscimol (Sigma-Aldrich catalog M1523) at 3.5 mM in saline were prepared, from which 0.4 ul (160 ng) was injected over 2 minutes using a World Precision Instruments syringe pump (model SP100i) with a 25 ul Hamilton syringe (model 702) while the mice were briefly anesthetized with Isofluorane (2-5% with O_2_). The drug was allowed to diffuse for 5 minutes after infusion before removing the cannula injector. Bilateral injections were performed serially, requiring the mouse to be under Isoflurane for a total of 14 minutes. After infusion, the mice were given 10 minutes to awake from anesthesia in their home-cage, after which they were placed on the T-maze to begin pairing.

Cannulated mice underwent daily pairing sessions until they met our criteria for being Learners (Sec. 2.6). On these initial pairing days, 0.4 ul of saline was infused bilaterally into auditory cortex before each session. After the subject mice met our criteria for being Learners, they had two additional days of pairing sessions where they received bilateral infusions of either muscimol or saline with the experimenter blind to which was being infused on which day.

## 3. Results

### 3.1. Initial pup search follows a location-based strategy

We sought to test whether female mice could use novel sensory cues to improve their search strategy for retrieving pups (Fig. 1). In our T-maze paradigm, each trial began with a subject mouse in its nest at the base of the T. One arm of the T (pseudo-randomized on each trial) was assigned as the arm to which a pup would be delivered when the subject entered that arm. Upon delivery, the pup would then be retrieved back to the nest.

During the initial session on the T-maze, there was considerable variability in how motivated different animals were in retrieving pups, as reflected in how many trials were completed in the 50 minute session (Fig. 2 bottom left). Nevertheless, 100% of the mice tended to select an arm to enter first on a given trial that was based on the arm from which they received a pup on the previous trial. We quantified this tendency by scoring each trial as a Correct or Wrong “location-based” choice based on entrance into the same or opposite arm, respectively, from which they had just been given a pup to retrieve (Fig. 2 top table). However, due to the pseudo-randomization, adherence to this “location-based” strategy was inefficient, and frequently resulted in the mice having to alternate arms before the pup would be given (Fig. 1 I & III).

**Figure 2:**
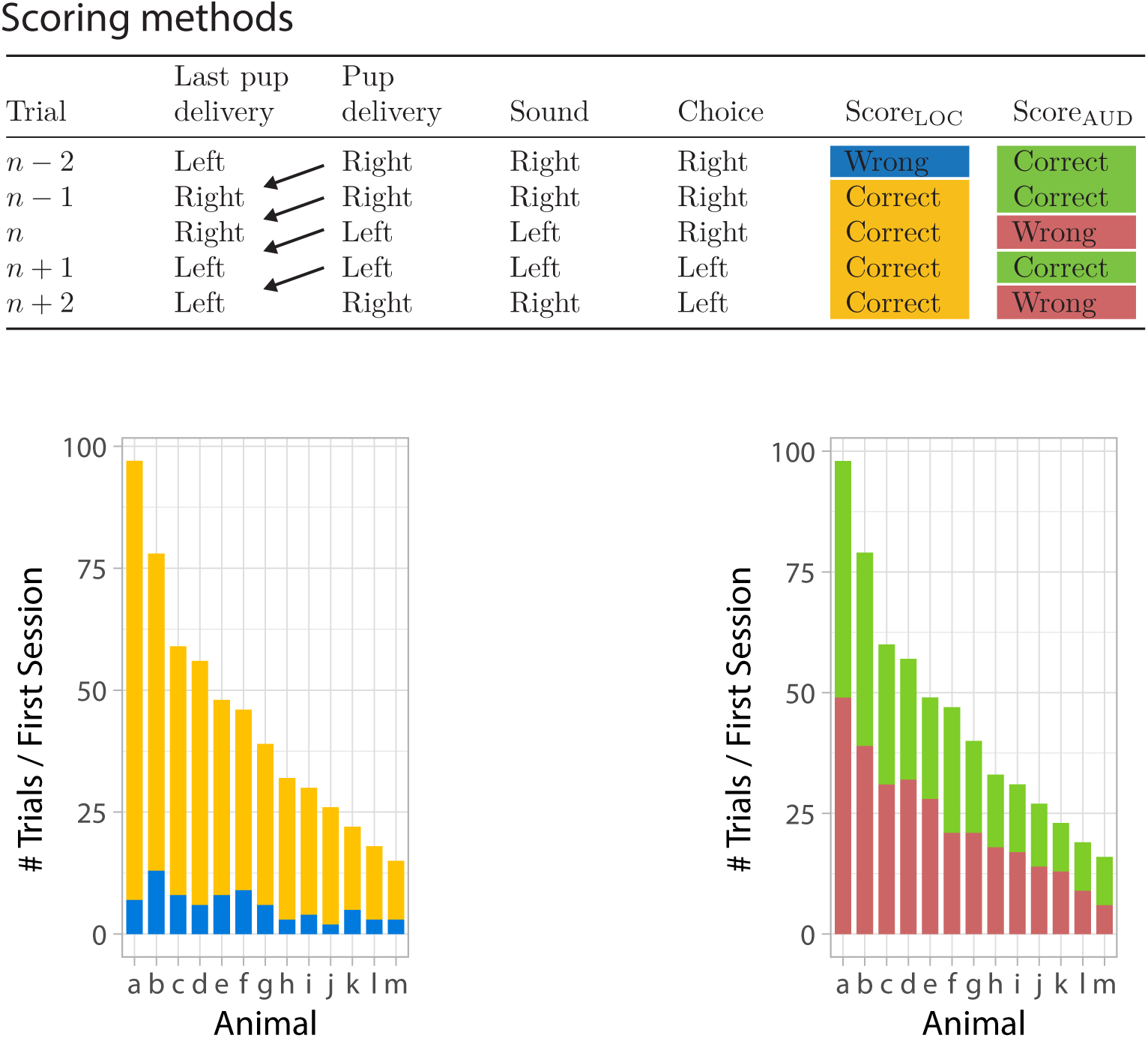
Behavioral strategies. (Top table) The sound- and location-based methods used to score pup search strategy. For the location-based strategy, Score_LOC_ was assigned a value of Correct when the subject chose the arm from which a pup was delivered on the previous trial, otherwise Score_LOC_ was assigned a value of Wrong. For the sound-based strategy, Score_SND_ was assigned a value of Correct when the subject chose the arm from which sound was being played on the current trial, otherwise Score_SND_ was assigned a value of Wrong. (Bottom left) On day 1, different virgin mice varied greatly in how many trials they completed during the 50 minutes of pairing; none reached the cap of 100 trials. All mice showed a propensity to use a location-based strategy to search for pups. (Bottom right) Performance according to a sound-based strategy was close to 50% for all mice, as expected if they did not recognize that the cue is associated with the arm for pup delivery.

Importantly, on each trial, a synthetic sound stream (see Materials and Methods) was also played back from a speaker at the end of the arm assigned for pup delivery. Hence, selecting an arm based on the cued sound could allow for a more efficient pup search strategy. We quantified the extent to which the mice utilized this auditory cue by scoring each trial as a Correct or Wrong “sound-based” choice based on entrance into the same or opposite arm, respectively, from which the sound was playing. During the initial session on the T-maze, each mouse’s performance according to this “sound-based” strategy was near chance, indicating that mice did not use the auditory cue on their first day in the T-maze (Fig. 2 bottom right).

### 3.2. Mice learn to use a sound-based search strategy by 7 days of pairing

After several daily sessions retrieving pups on the maze, mice learned to shift from using the location-based to the sound-based strategy. For example, on its first day in the T-maze, the pup-naive, virgin mouse in Fig. 3A chose arms based on where it previously received a pup in more than 90% of trials. Between days 5 and 6 though, the number of trials scored Correct based on location decreased considerably, while those scored Correct based on sound increased (Fig. 3B). This trend continued over subsequent days until the mouse’s location-based performance dropped to chance, and its sound-based performance reached nearly 90% (Fig. 3C & D). Meanwhile, the total number of trials completed each day increased to the cap of 100 trials in 50 minutes (Fig. 3A or B) as the animal reached its decision point faster (i.e. higher reciprocal latency, Fig. 3E).

**Figure 3:**
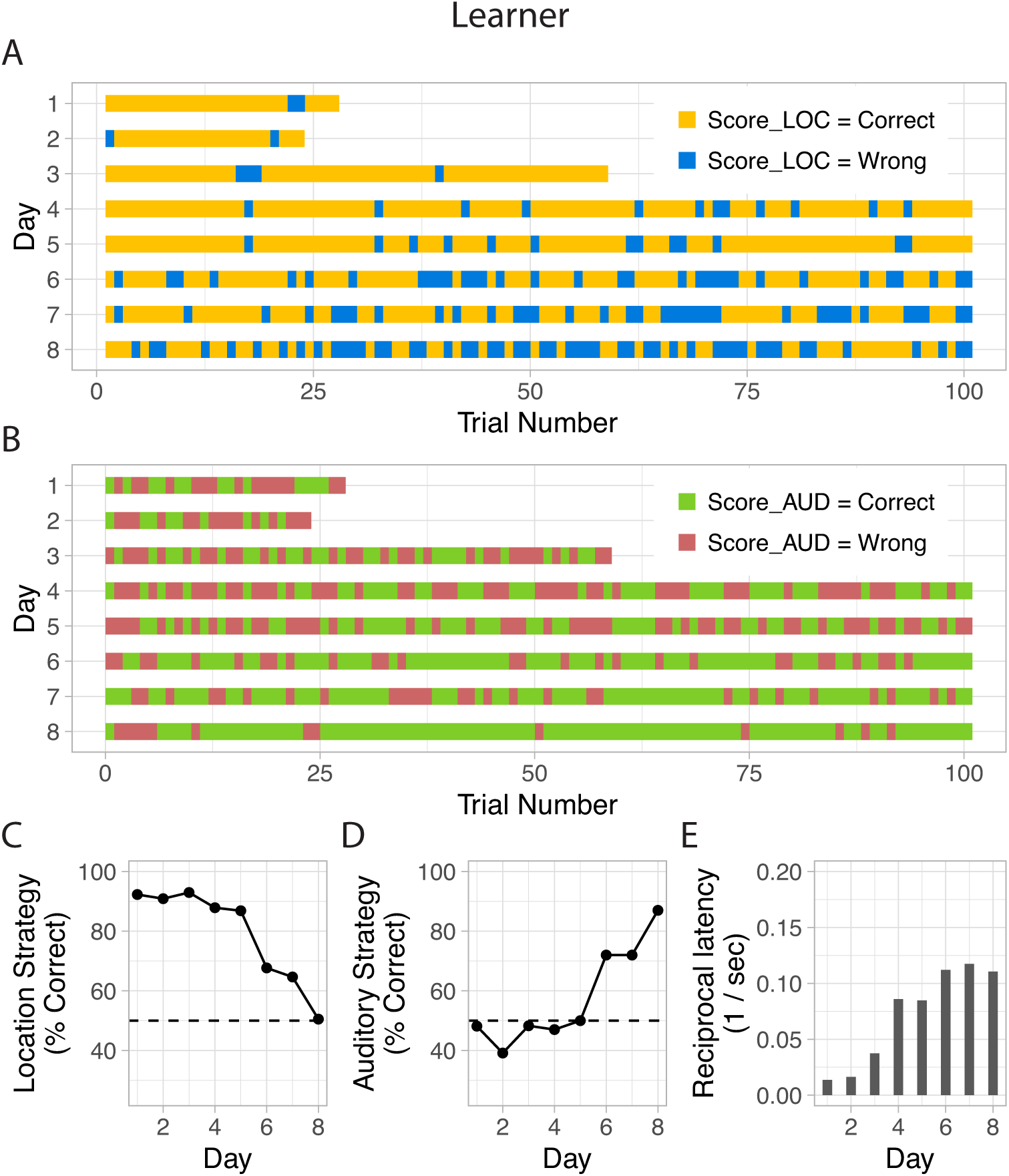
Example Learner mouse that shifted its retrieval strategy over sessions. (A) Trials scored according to the location-based strategy. (B) Same trials scored according to the sound-based strategy. (C) Daily percent correct score values for the location-based method of scoring. Dashed line represents chance at 50%. (D) Daily percent correct score values for the sound-based method of scoring. (E) Daily reciprocal latency values.

Across our sample of naive, virgin mice, we found that after 7 days of pairing, 9 mice (69%) performed significantly better than chance according to the sound-based strategy, averaging 72% (standard error (SE) = 3%) on day 7. On day 8, we tested these “Learners” with a second example of the sound stream, with acoustic properties matched to those of the originally paired stream (see Materials and Methods). Sound-based performance remained high (“Sound on” trials in Fig. 4A), ruling out potential pseudo-replication concerns with our pairing stimulus. We also introduced silent catch trials on day 8, where on 20% of trials, no sound cue was delivered from the selected arm but the mice were still rewarded with a pup upon entry. Average sound-based performance was 77% (SE = 4%) during “Sound on” playback trials and 47% (SE = 5%) for silent, “Sound off” trials (Fig. 4A). This performance drop was not attributable to how motivated the animals were to quickly reach a pup, since reciprocal latency (higher values indicate faster choices) did not differ between the two conditions (Fig 4B). Moreover, at 60% (SE = 5%), reliance on a location-based strategy during these randomly interleaved “Sound off” trials did not increase much above chance levels, suggesting that these animals did not revert to the less efficient strategy despite the intermittent loss of the sound cue. Taken together, these results suggest that Learners had indeed learned to use the sound to improve their strategy for efficiently choosing an arm to retrieve a pup.

### 3.3. Auditory learning of pup-predictive sounds is separable from motivation to retrieve pups

Across pairing days, we saw gradual increases in both the sound-based performance and the speed with which animals made choices (Fig 4). Therefore we next asked about the relationship between these variables, which served as proxies for sensory learning and motivation, respectively. Across individuals, those mice that learned by days 7 and 8 to perform better using the sound cue were also retrieving faster, a correlation that was significant (Fig. 5A). However, when looking within animals, the trial-by-trial time to make a decision was not significantly different for trials scored as Correct or Wrong according to either strategy (Fig. 5B & C). Hence, even though highly motivated animals tended to use the sound cue more successfully, the factors determining fluctuations in decision latency on a trial-by-trial basis were different from those that dictated which arm was chosen. Importantly, the T-maze paradigm allowed us to separately assay sound-based versus motivation-driven components of pup retrieval.

**Figure 4:**
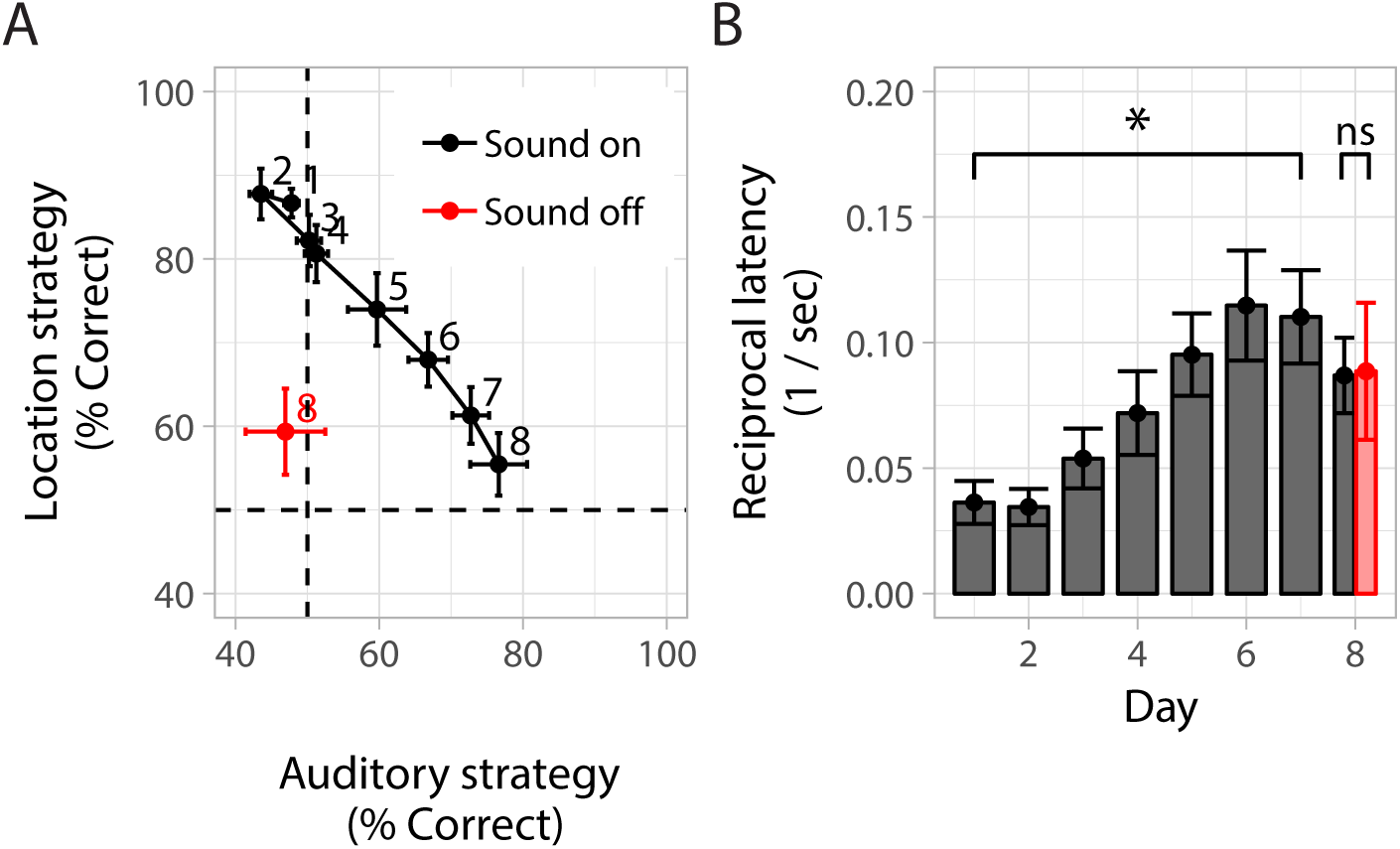
Group data showing auditory learning in virgin mice. (A) Initially, female mice used a location-based strategy, where they chose to enter the arm that they last retrieved a pup from. Reliance on this strategy decreased over pairing as the mice developed a new sound-based strategy, where they chose to enter the arm indicated by sound playback. Performance as measured by the sound-based strategy was significantly different on silent trials (*p* < 0.05), confirming the mice were using the sound cue. (B) As Learners shifted their strategy, their motivation to retrieve significantly increased (*p* < 0.05). Motivation to retrieve was not significantly different sound on/off (gray/red right bars) trials.

**Figure 5:**
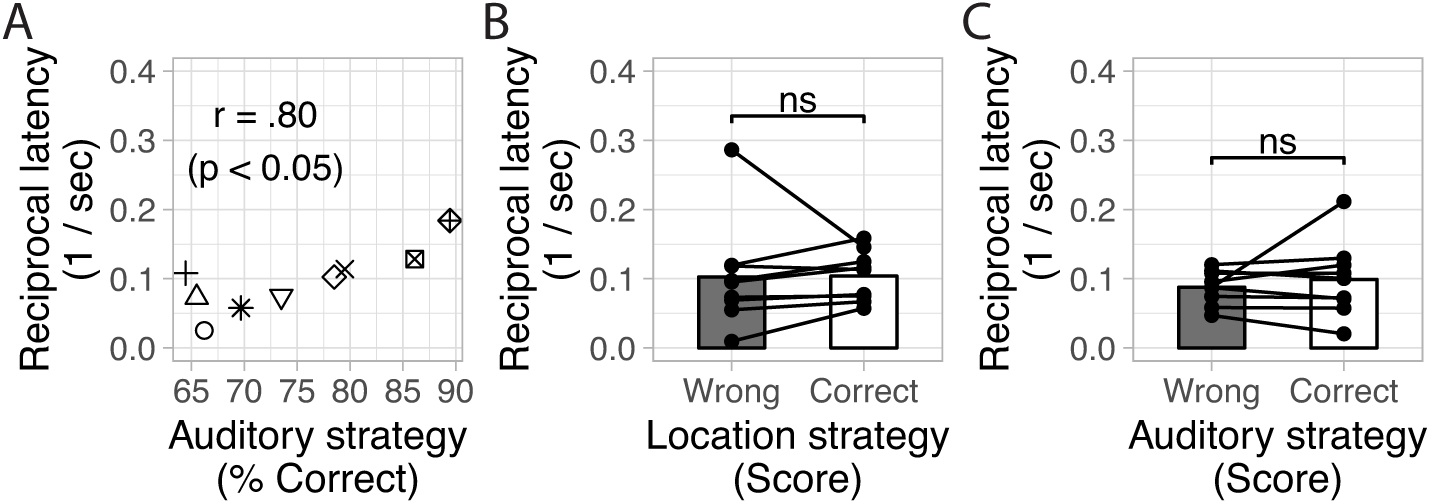
Relationship between motivation and strategy, measured from “Sound on” trials pooled over days 7 & 8. (A) For mice that learned, reciprocal latency and sound-based strategy percent correct were correlated across animals. (B) However, when looking within animals, their latency to make a choice was not significantly different depending on whether or not they got a trial correct according to the sound-based strategy. (C) Similarly, their latency to make a choice was not significantly different depending on whether or not they got a trial correct according to the location-based strategy.

### 3.4. Learners form a sound-specific sensory memory associated with pups

Since Learners came to use the sound to guide their search, we investigated whether their improvement coincided with forming a long-term sensory memory. Using a separate group of 4 naive mice paired over 6 consecutive days, we assayed learning retention 10-20 days after their last day of training. We assessed performance over just the first 10 trials of a session to minimize the effect of renewed reinforcement. Performance on the last pairing day was significantly above chance, as expected for Learners (Fig. 6A). More importantly, when these animals were tested for retention at a remote time point, all continued to perform above chance. This result demonstrates that the sound’s learned meaning persisted beyond one week, consistent with a long-term memory.

**Figure 6:**
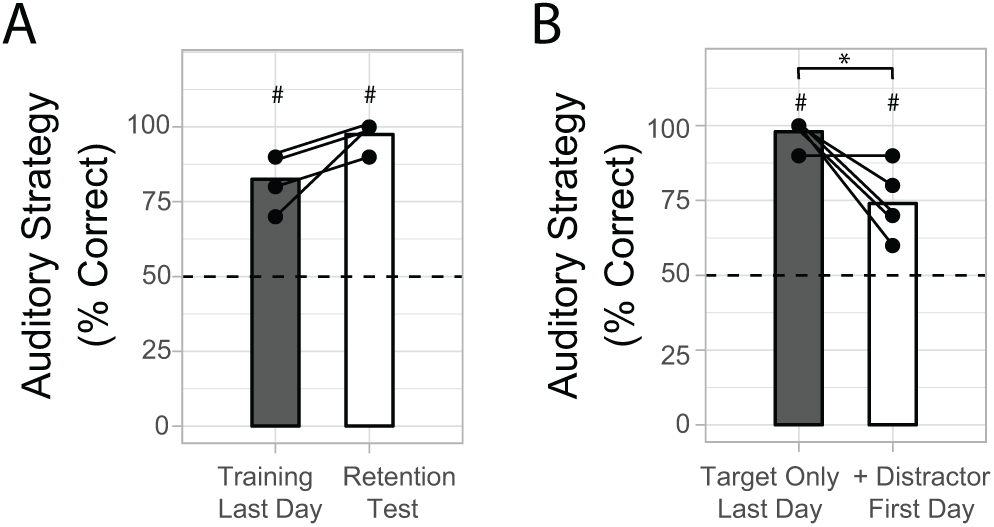
Cue-specific sensory memory. (A) Mice trained on the T-maze correctly preferred the sound arm during the first 10 trials on both their last day of regular pairing, and the retention test 10-20 days later (#: *p* < 0.05, z-test of proportion vs. 50% chance). (B) Learners displayed a small but significant drop in performance over the first 10 trials after a distractor sound was played from the non-target speaker (*: *p* < 0.05, paired t-test), suggesting they perceived the new sound. Nevertheless, they retained a preference for the pup-predictive target sound even over these few trials (#: *p* < 0.05, z-test of proportion vs. 50% chance).

To gain further insight into what animals had learned, we next asked whether Learners discriminated their target sound from other sounds, or simply learned to approach a speaker playing any sound. We paired the target sound presented by itself (as before) with pups for another group of 5 mice in the T-maze. All Learners performed above chance on their last day of pairing with just the target (Fig. 6B). The next day, we introduced from the first trial an arbitrary distractor sound played as a non-synchronized stream from the non-target speaker (see Materials and Methods). Performance dropped slightly but significantly over the first 10 trials in the session, yet remained significantly above chance. These results suggest that Learners continued to recognize and prefer the target sound, even though they perceived the novel distractor sound.

Finally, we reasoned that if the sound itself was recognized as a localization cue for retrieving pups, then Learners should be able to generalize their learning to a novel context. Two additional naive mice were paired on the T-maze (70% and 90% on first 10 trials on the last day of T-maze training), and then tested in a rat-sized cage partitioned into a Y-maze (see Materials and Methods). Their performance over their very first 10 trials in the new context remained far above chance (90% and 80%, respectively). As a comparison, five animals that had not previously been paired in either the T-maze or Y-maze performed at chance in their very first 10 trials in the Y-maze (46 *±* 5%). The fact that performance easily generalized to a novel choice context supports the notion that T-maze animals had learned the meaning of the sensory cue rather than just the operant context.

### 3.5. Learned search strategy, but not motivation, depends on auditory cortical processing

At a neural level, the auditory cortex has been implicated previously in pup vocalization-associated neural plasticity, and we hypothesized that it would also play a key role in the online usage of the sound-based strategy for efficient pup retrieval. To test this hypothesis, we bilaterally infused muscimol into auditory cortex (see Materials and Methods) in Learners.

We cannulated and paired 9 mice up to criteria. We found (Fig. 7A & B) that bilateral infusion of the GABA agonist muscimol into auditory cortex after animals reached criteria significantly decreased performance as measured by the sound-based strategy, while also significantly improving performance as measured by the location-based strategy. Hence, silencing auditory cortex shifted the strategy mice used from a sound-based back towards a location-based one. On the other hand, we found (Fig. 7C) no significant effect of treatment on trial reciprocal latency, indicating that auditory cortical inactivation did not strongly influence the time taken for mice to leave the nest and choose an arm.

**Figure 7:**
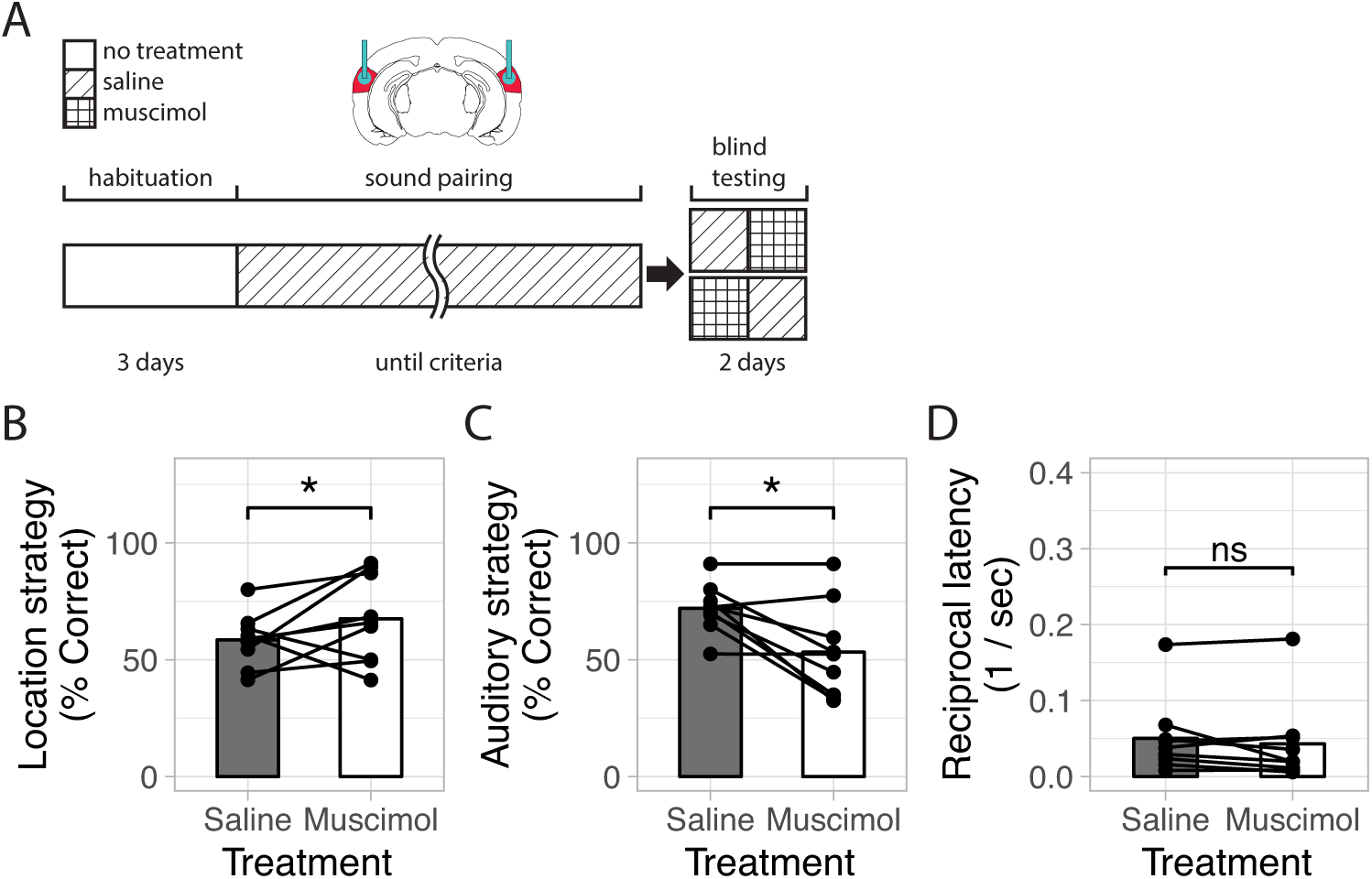
Improved strategy, but not retrieval, depends on sensory cortical processing. (A) Auditory cortex was bilaterally cannulated (bregma = −3.04 mm, midline = ± 3.75 mm, z = −0.5 mm from cortical surface). On pairing days, saline was infused 24 minutes prior to pairing. On testing days, either saline or muscimol (counterbalanced) was infused 24 minutes prior to conditioning. (B) Silencing auditory cortical activity significantly decreases performance when measured by the sound-based strategy (*p* < .05). (C) Silencing auditory cortical activity significantly increases performance when measured by the location-based strategy (*p* < .05). (D) Silencing auditory cortical activity does not significantly alter reciprocal latency of retrieval approach.

### 3.6. Maternal state increases motivation to retrieve and accelerates change in strategy

Next, at a physiological level, we hypothesized that the maternal state could play a role in facilitating associative learning for auditory cues [19]. To test this hypothesis, we compared lactating mothers and cocaring females in the T-maze paradigm. Cocarers are virgin sisters of a mother, and they were housed with the mother (and her litter) from before parturition through T-maze pairing. Thus, both animal groups had experience caring for pups, but mothers underwent the additional physiological and hormonal changes associated with pregnancy, birth, and lactation. We paired 13 lactating mothers and 12 cocarers in our paradigm. Of these, 11 (84%) of the lactating mothers and 6 (50%) of the cocarers met our criteria for having learned the task. Our analyses were restricted to these subset of mice.

Over pairing, the reciprocal latency for mothers to approach and enter one of the arms on the T-maze increased faster than for cocarers (Fig. 8A). We found a significant effect of animal group and a significant interaction between animal group and testing day. For mothers, the greatest changes in reciprocal latency occurred over the first three days. This effect was not seen for cocaring mice, whose reciprocal latency did not significantly change with pairing, which demonstrates that cocarers were able to learn without an increase in our proxy for maternal motivation. On days 2-8 we found a significant effect of animal group on trial reciprocal latency.

**Figure 8:**
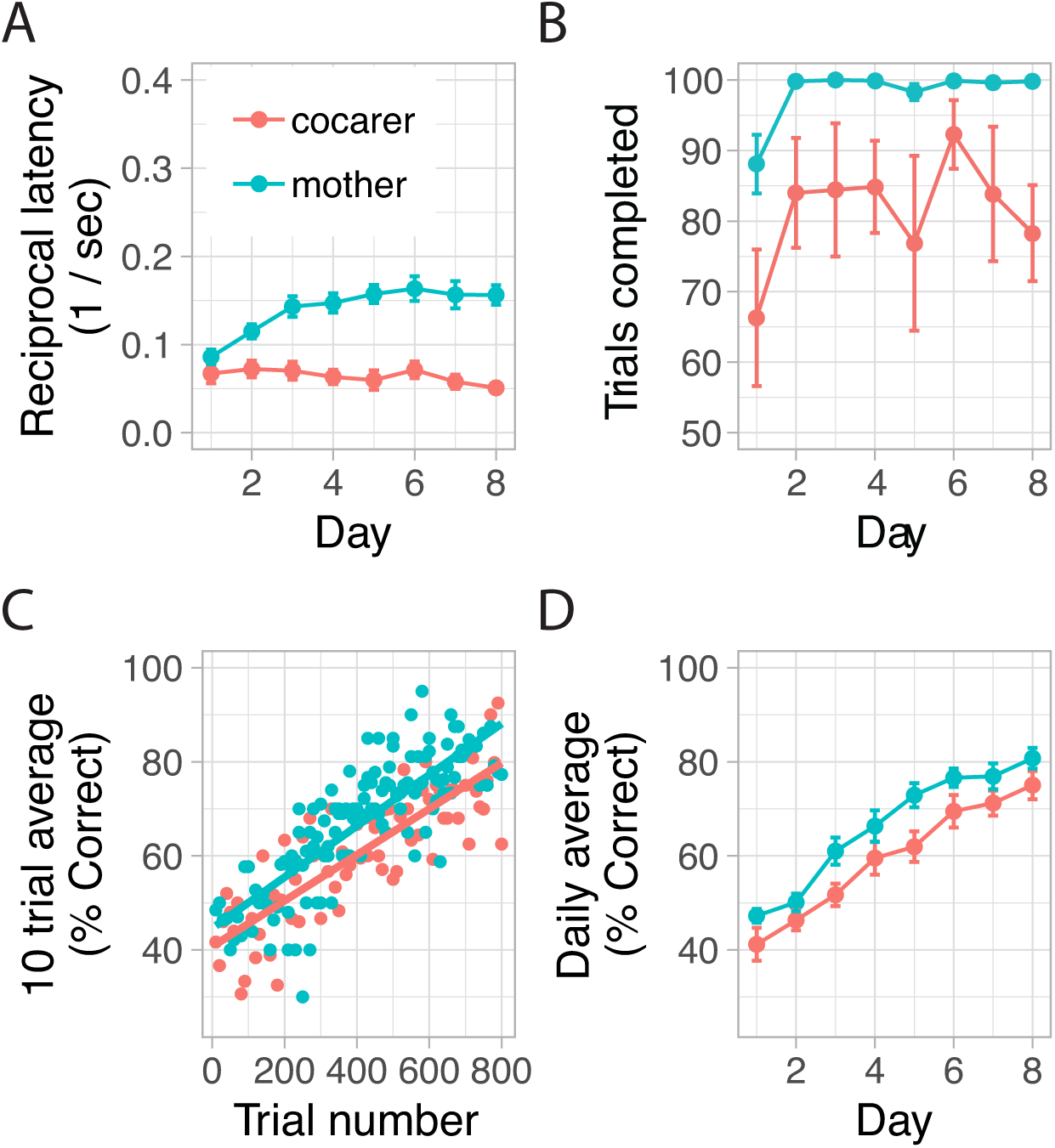
Maternal state accelerates change in strategy by increasing motivation to retrieve. (A) Reciprocal latency increases faster for mother mice than cocarers over days (*p* < .05). (B) Mother mice complete more trials per day than cocarers. (C) On a per trial basis, mother mice have a significantly higher offset in performance when measured by the sound-based strategy (*p* < .05). (D) Combined, motherhood allows mice to improve performance according to the sound-based strategy faster than cocarers over days (*p* < .05).

A consequence of the mothers having higher motivation to search for and retrieve pups was that mothers completed more trials per day than cocarers (Fig. 8B). To visualize learning after matching for the number of trials completed (rather than number of days), we filtered the sound-based trial scores for each animal (10 trial running averages), and fit regression lines to these points (Fig. 8C). Even on this basis, mothers showed enhanced learning. We quantified this by fitting a model to the unfiltered sound-based trial scores as a function of trial number to confirm a significant effect of animal group. Therefore, motherhood enhanced learning on a per trial basis in addition to increasing the trials completed per day. When combined together, these factors allowed mothers to adapt their approach strategy in response to the non-ethological but pup-predictive auditory cue faster over days. Indeed, sound-based performance over days (Fig. 8D) was significantly higher in mothers.

## 4. Discussion

Here we showed that with infant experience, sensory learning of infant-associated cues can come to improve a stereotypical maternal behavior in female mice, irrespective of changes in maternal motivation. Adult females search for and retrieve pups, which are intrinsically rewarding, and this behavior can be used to reinforce learning novel cues that are predictive of where to find pups. We repetitively delivered pups into the arms of a T-maze to encourage search behavior. The default strategy mice used to select an arm was based on the spatial memory for where they were last given a pup to retrieve (Fig. 1-2). However, mice then learned to use a novel pup-associated sound to instead guide more efficient search and approach (Fig. 3-4). The motivation to retrieve pups also increased with experience, but motivation alone could not explain how well the new sound cue was used to guide pup search (Fig. 5). This altered strategy utilized a long-term memory of the specific, pup-predictive sound (Fig. 6), and required auditory cortical activity to execute (Fig. 7). Finally, motherhood accelerated how quickly this new strategy was acquired over days (Fig. 8). Below we discuss these results in the context of prior work on maternal behavior and sensory learning.

Maternal behavior is thought to be governed largely by neural circuits that are shaped by evolution and “pre-wired” to rapidly support maternal responsiveness once engaged. A recent study demonstrated that the circuitry for parental behavior is present in both females and males, and can be effectively “switched on” through optogenetics so that even virgin male mice, which would normally be hostile towards pups, abruptly become more caring [20]. Under normal conditions, maternal hormones are believed to modulate how such circuits enable a transition from being neglectful towards pups to being a responsive mother [3]. Notably though, many studies of maternal behavior in rodents rely primarily on assessing just the presence or absence of discrete behaviors like crouching over pups or pup retrieval [2]. While this can be done quantitatively, such measurements typically do not capture *how* maternal behaviors are being performed, leaving open questions about the strategies for providing infant care, and whether experience may alter these.

Importantly, interactions with infants can be socially rewarding, and thus infant reinforcement may shape how an animal learns to improve maternal care over time - a topic of potential clinical relevance [21]. We used female rodents, for which pups act as a reinforcer [22, 23], and pup retrieval, which is a natural maternal behavior that depends on the integrity of the mesocorticolimbic dopaminergic system [24], an essential part of learning and goal-directed behaviors. Our use of a T-maze paradigm for pup search and retrieval in mice allowed us to assess whether pup reinforcement could drive the more efficient use of sensory cues that are predictive of where to find a pup. With this paradigm, we could study *how* new information is being incorporated to change a stereotypical maternal behavior. In particular, we could track how animals switch from using a location-based search strategy to using sounds to find pups, and the extent to which incorporating this new information is influenced by maternal motivation. Hence, our paradigm opens the door to rigorously studying a more cognitive aspect of parental behavior in rodent models.

Our findings suggest that infant search strategy is regulated separately from maternal motivation. Mice varied considerably on day 1 in their motivation to retrieve pups, yet they uniformly applied a location-based search strategy in the T-maze at the start of training (Fig. 2 bottom left). Furthermore, day 1 cocarers had similar average levels of maternal motivation (Fig. 8A) as day 6-8 naives (Fig. 4B) - as expected after pup exposure in the home cage and/or T-maze [25] - yet that was insufficient to guarantee similar sound-based performance. In fact, even as cocarers improved in their use of a sound-based strategy over days, their reciprocal latency remained constant, further suggesting that motivation and performance are decouplable characteristics. Most tellingly was that on a trial-by-trial basis, whether animals were Correct or Wrong in their arm choice (Fig. 5B-C), did not change their latencies to choose an arm. Additionally, latencies were not different on the final day between trials with or without sound (Fig. 4B), demonstrating that motivation can remain high even without the ability to apply the sound-based strategy. Hence, even though more maternally motivated animals can eventually reach higher levels of performance (Fig. 5A), the ability to exploit pup-predictive cues for search is not simply a monotonic function of one’s motivation to engage with pups. This is important to factor into the interpretation of studies that only use the amount of pup retrieval as a proxy for sensory learning about pup cues without directly testing preferred approach to the learned sound.

Our results further suggest that maternal physiological state impacts both motivation and performance in a sensory learning task. The motivation to search for and retrieve pups reached its highest levels for mothers compared to the other animal groups, a result consistent with prior work on hormonal contributions to maternal responsiveness in rodents [26]. Notably, for both animal groups, the pups used during training were foster pups, so the enhancement by motherhood could not be explained by extra attention to one’s own pups. Moreover, compared to cocarers, mothers switched from the location-based to the sound-based strategy the fastest, even on a trial-matched basis (Fig. 8C). This was not obvious *a priori* since the location-based strategy to return to the last location a pup was found requires spatial memory, which prior studies suggest can be reinforced by motherhood [27]. Hence, motherhood heightens learning the most efficient strategies to reach pups to retrieve.

Our conclusions depended on being able to reliably assay and quickly track the acquisition of the sound-based search strategy. Several aspects of the T-maze paradigm are critical for this success. First, subject mice do not have to be trained to perform a new behavior - such as lever pressing or nose poking [23, 28] - in order to demonstrate their recognition of the sound’s behavioral relevance. By minimizing the motor shaping steps that are common in operant paradigms, we reduce the time required to conduct experiments, increasing throughput. In fact, since subjects normally retrieve pups back to the nest, the behavior naturally sets animals up to quickly begin their next trial. Moreover, because we tap into the ethological behavior, animals have a baseline strategy for completing the task that they apply from the very beginning of training. Our ability to quantify that strategy allows us to track the transition between intrinsic and learned strategies in a way that escapes investigation in most operant paradigms.

Second, because we continually deliver pups to reinforce learning to use the non-ethological sound to guide search, individual animals can perform nearly endless trials without habituating - a useful feature for future neurophysiological studies during behavior. This is also an advantage for assessing individual recognition and learning, compared to previous studies of USV recognition [29] that relied on group data, since individual mice could only conduct up to 6 trials before habituating to sounds that were not reinforced with actual pups. Introducing “catch-trials” without sounds helps ensure that this performance is actually based on recognizing the auditory cue. Finally, our test using a distractor sound demonstrates that after training, subjects discriminate the learned target from other acoustic categories (even in the same frequency range), suggesting a relatively specific sound memory.

To help elucidate the neural mechanisms by which the new sound was being used to guide search, we showed that auditory cortical activity was necessary to express the learned, sound-based strategy (Fig. 7). We targeted auditory cortex because it is the first site along the auditory pathway where neural activity is correlated with perception [30], and its plasticity is thought to underlie memory formation [31]. Auditory cortical activity also supports the ability to localize sound sources [32], although that function may be less relevant for our T-maze paradigm. Our pup-predictive sound was presented from the left or right side at the decision point, and prior studies suggest that auditory cortical activity is not needed to localize the hemisphere from which a long duration sound is played [33, 34]. On the other hand, the necessity of auditory cortical activity for accurately choosing an arm to search, despite other auditory pathways for influencing behavioral responses to sounds [35], is consistent with auditory cortical projections affecting behavioral choices in sound-based tasks [36, 37]. Our results lay the groundwork for future studies using the T-maze to dissect the circuitry involved in socially reinforced auditory learning.

Finally, given that we can now robustly demonstrate sensory associative learning in this social context, future studies can investigate whether there is neural plasticity within auditory cortex, or other sites along the auditory pathway, for the paired sound. Of particular interest would be to compare any observed plasticity to changes reported for USVs themselves [4, 5, 6, 8, 38, 39], which are the ethological cue that elicits search and retrieval by mothers. Those neural changes have been hypothesized to support the association between pups and USVs after the subject has had experience with vocalizing pups [14]. An alternative, albeit not mutually exclusive, hypothesis is that USV responses are unlocked by the unique neurochemical and hormonal conditions present during motherhood [40]. These two hypotheses make different predictions about how much experience is required for mothers to display preferred approach to USVs versus non-ethological sounds. By comparing how the maternal state influences the acquisition of preferred approach for ethological versus non-ethological sounds, the T-maze paradigm could be used to validate a sensory associative learning component for the acquisition of preferred approach to USVs during natural maternal behavior.

## 5. Acknowledgements

This work was supported by the National Institutes of Health Grants R01-DC-8343 (RCL) and T32-HD-071845 (KKC) and Emory University’s Laney Graduate School Summer Opportunity for Academic Research program (BBT). Contributions: AGD and RCL designed research; AGD, CB, LMP, KKC, DBK and BBT carried out experiments; HW and KL contributed to methods; AGD, CB, LMP and DBK analyzed data; AGD and RCL wrote the paper. We thank Vanessa Wong for experimental assistance.

## References

[1] G. Ehret, Infant rodent ultrasounds–a gate to the understanding of sound communication, Behav. Genet. 35 (1) (2005) 19–29.

[2] M. Numan, T. R. Insel, The Neurobiology of Parental Behavior, Springer Science & Business Media, 2006.

[3] J. Kohl, A. E. Autry, C. Dulac, The neurobiology of parenting: A neural circuit perspective, Bioessays 39 (1) (2017) 1–11.

[4] R. C. Liu, J. F. Linden, C. E. Schreiner, Improved cortical entrainment to infant communication calls in mothers compared with virgin mice, Eur. J. Neurosci. 23 (11) (2006) 3087–3097.

[5] R. C. Liu, C. E. Schreiner, Auditory cortical detection and discrimination correlates with communicative significance, PLoS Biol. 5 (7) (2007) e173.

[6] E. E. Galindo-Leon, F. G. Lin, R. C. Liu, Inhibitory plasticity in a lateral band improves cortical detection of natural vocalizations, Neuron 62 (5) (2009) 705–716.

[7] L. Cohen, G. Rothschild, A. Mizrahi, Multisensory integration of natural odors and sounds in the auditory cortex, Neuron 72 (2) (2011) 357–369.

[8] B. J. Marlin, M. Mitre, J. A. D’amour, M. V. Chao, R. C. Froemke, Oxytocin enables maternal behaviour by balancing cortical inhibition, Nature 520 (7548) (2015) 499–504.

[9] M. Mitre, B. J. Marlin, J. K. Schiavo, E. Morina, S. E. Norden, T. A. Hackett, C. J. Aoki, M. V. Chao, R. C. Froemke, A distributed network for social cognition enriched for oxytocin receptors, J. Neurosci. 36 (8) (2016) 2517–2535.

[10] F. W. Ohl, H. Scheich, W. J. Freeman, Change in pattern of ongoing cortical activity with auditory category learning, Nature 412 (6848) (2001) 733–736.

[11] N. M. Weinberger, Specific long-term memory traces in primary auditory cortex, Nat. Rev. Neurosci. 5 (4) (2004) 279–290.

[12] M. L. Caras, D. H. Sanes, Top-down modulation of sensory cortex gates perceptual learning, Proc. Natl. Acad. Sci. U. S. A. 114 (37) (2017) 9972–9977.

[13] S. Bennur, J. Tsunada, Y. E. Cohen, R. C. Liu, Understanding the neurophysiological basis of auditory abilities for social communication: a perspective on the value of ethological paradigms, Hear. Res. 305 (2013) 3–9.

[14] S. B. Banerjee, R. C. Liu, Storing maternal memories: hypothesizing an interaction of experience and estrogen on sensory cortical plasticity to learn infant cues, Front. Neuroendocrinol. 34 (4) (2013) 300–314.

[15] J. M. Stern, D. A. Mackinnon, Postpartum, hormonal, and nonhormonal induction of maternal behavior in rats: effects on t-maze retrieval of pups, Horm. Behav. 7 (3) (1976) 305–316.

[16] D. S. Stolzenberg, E. F. Rissman, Oestrogen-independent, experience-induced maternal behaviour in female mice, J. Neuroendocrinol. 23 (4) (2011) 345–354.

[17] R Core Team, R: A Language and Environment for Statistical Computing, R Foundation for Statistical Computing, Vienna, Austria (2017). URL https://www.R-project.org/

[18] D. Bates, M. Mächler, B. Bolker, S. Walker, Fitting linear mixed-effects models using lme4, Journal of Statistical Software 67 (1) (2015) 1–48. doi: 10.18637/jss.v067.i01.

[19] F. G. Lin, E. E. Galindo-Leon, T. N. Ivanova, R. C. Mappus, R. C. Liu, A role for maternal physiological state in preserving auditory cortical plasticity for salient infant calls, Neuroscience 247 (2013) 102–116.

[20] Z. Wu, A. E. Autry, J. F. Bergan, M. Watabe-Uchida, C. G. Dulac, Galanin neurons in the medial preoptic area govern parental behaviour, Nature 509 (7500) (2014) 325–330.

[21] T. Field, Postpartum depression effects on early interactions, parenting, and safety practices: a review, Infant Behav. Dev. 33 (1) (2010) 1–6.

[22] A. S. Fleming, M. Korsmit, M. Deller, Rat pups are potent reinforcers to the maternal animal: effects of experience, parity, hormones, and dopamine function, Psychobiology 22 (1) (1994) 44–53.

[23] A. Lee, S. Clancy, A. S. Fleming, Mother rats bar-press for pups: effects of lesions of the mpoa and limbic sites on maternal behavior and operant responding for pup-reinforcement, Behav. Brain Res. 108 (2) (2000) 215–231.

[24] S. Hansen, C. Harthon, E. Wallin, L. Löfberg, K. Svensson, Mesotelencephalic dopamine system and reproductive behavior in the female rat: effects of ventral tegmental 6-hydroxydopamine lesions on maternal and sexual responsiveness, Behav. Neurosci. 105 (4) (1991) 588.

[25] J. S. Rosenblatt, Nonhormonal basis of maternal behavior in the rat, Science 156 (3781) (1967) 1512–1514.

[26] J. S. Rosenblatt, A. Olufowobi, H. I. Siegel, Effects of pregnancy hormones on maternal responsiveness, responsiveness to estrogen stimulation of maternal behavior, and the lordosis response to estrogen stimulation, Horm. Behav. 33 (2) (1998) 104–114.

[27] C. H. Kinsley, L. Madonia, G. W. Gifford, K. Tureski, G. R. Griffin, C. Lowry, J. Williams, J. Collins, H. McLearie, K. G. Lambert, Motherhood improves learning and memory, Nature 402 (6758) (1999) 137–138.

[28] E. G. Neilans, D. P. Holfoth, K. E. Radziwon, C. V. Portfors, M. L. Dent, Discrimination of ultrasonic vocalizations by CBA/CaJ mice (mus musculus) is related to spectrotemporal dissimilarity of vocalizations, PLoS One 9 (1) (2014) e85405.

[29] G. Ehret, B. Haack, Ultrasound recognition in house mice: Key-Stimulus configuration and recognition mechanism, J. Comp. Physiol. 148 (2) (1982) 245–251.

[30] J. K. Bizley, Y. E. Cohen, The what, where and how of auditory-object perception, Nat. Rev. Neurosci. 14 (10) (2013) 693–707.

[31] K. N. Shepard, M. P. Kilgard, R. C. Liu, Experience-Dependent plasticity and auditory cortex, in: Neural Correlates of Auditory Cognition, Springer Handbook of Auditory Research, Springer New York, 2013, pp. 293–327.

[32] A. J. King, V. M. Bajo, J. K. Bizley, R. A. A. Campbell, F. R. Nodal, A. L. Schulz, J. W. H. Schnupp, Physiological and behavioral studies of spatial coding in the auditory cortex, Hear. Res. 229 (1-2) (2007) 106–115.

[33] S. Malhotra, S. G. Lomber, Sound localization during homotopic and heterotopic bilateral cooling deactivation of primary and nonprimary auditory cortical areas in the cat, J. Neurophysiol. 97 (1) (2007) 26–43.

[34] F. R. Nodal, O. Kacelnik, V. M. Bajo, J. K. Bizley, D. R. Moore, A. J. King, Lesions of the auditory cortex impair azimuthal sound localization and its recalibration in ferrets, J. Neurophysiol. 103 (3) (2010) 1209–1225.

[35] G.-W. Zhang, W.-J. Sun, B. Zingg, L. Shen, J. He, Y. Xiong, H. W. Tao, L. I. Zhang, A non-canonical Reticular-Limbic central auditory pathway via medial septum contributes to fear conditioning, Neuron 97 (2) (2018) 406–417.e4.

[36] L. Guo, J. T. Weems, W. I. Walker, A. Levichev, S. Jaramillo, Choice-Selective neurons in the auditory cortex and in its striatal target encode reward expectation, J. Neurosci. 39 (19) (2019) 3687–3697.

[37] Q. Xiong, P. Znamenskiy, A. M. Zador, Selective corticostriatal plasticity during acquisition of an auditory discrimination task, Nature 521 (7552) (2015) 348–351.

[38] K. Krishnan, B. Y. B. Lau, G. Ewall, Z. J. Huang, S. D. Shea, MECP2 regulates cortical plasticity underlying a learned behaviour in adult female mice, Nat. Commun. 8 (2017) 14077.

[39] G.-I. Tasaka, C. J. Guenthner, A. Shalev, O. Gilday, L. Luo, A. Mizrahi, Genetic tagging of active neurons in auditory cortex reveals maternal plasticity of coding ultrasonic vocalizations, Nat. Commun. 9 (1) (2018) 871.

[40] C. I. Bargmann, Beyond the connectome: how neuromodulators shape neural circuits, Bioessays 34 (6) (2012) 458–465.

